# Point set registration for combining fluorescence microscopy methods

**DOI:** 10.1101/2024.06.22.600172

**Authors:** Ivo Severins, Chirlmin Joo, John van Noort

## Abstract

**SUMMARY:** Implementation of combined microscopy methods provides valuable information across various scientific applications. However, aligning the datasets and finding the correct point correspondence poses a challenge, especially for large, randomly distributed point sets that are subject to positional errors and missing points. Here, we provide a three-step procedure to perform point set registration, which can be applied to datasets with millions of points and stays robust even when only 10% of the points correspond. In the first global step, the scaling and rotation parameters for the imaging systems are determined once on a smaller calibration dataset using a geometric hashing algorithm. When the global transformation is known, full experimental datasets can be registered by performing step two: a course registration using cross-correlation, and step three: a precise registration to fine-tune the transformation. After these three steps, point correspondence is determined by setting a distance threshold based on a statistical model of random point sets that additionally provides the matching error. We have demonstrated its successful implementation in coupling fluorescence and sequencing methodologies. To enable wide application of these point set registration and correspondence algorithms we provide a python library called MatchPoint.

## INTRODUCTION

Looking at a problem from a different perspective has always helped to obtain new insights. In microscopy this is no different; combining observations from different imaging methods can provide information that no method could provide alone. An example is the combination of images from light and electron microscopy (Mohammadian et al. 2019) allowing molecular localization using fluorescence and providing structural information with electron microscopy. Another example is the combination of phase contrast and immunofluorescence images, the former following the lineages of dividing cells while the latter determines cellular expression of a marker protein (Becker et al. 2013). Our own application combines next-generation Illumina sequencing with single-molecule fluorescence microscopy allowing us to perform single-molecule experiments for millions of different DNA sequences simultaneously (Severins et al. in review).

While the difference between the two combined microscopy methods provides its value, it comes with the challenge of aligning the resulting images and datasets. Registering images directly may be difficult as the features used for alignment may have different appearances for the two microscopy methods. To overcome this, registration is instead performed on the coordinates of the features that are located in the image. Although this takes away one obstacle, many others remain. In the case of registering point sets for combining single-molecule measurements with sequencing, the challenges originate from: (1) the unknown imaging parameters of the sequencer and the output coordinates given in arbitrary units; (2) the randomly distributed positions of molecules and sequencing clusters which exclude the use of shape detection approaches; (3) a large difference in the scale of the images, i.e. roughly 400 single-molecule images fit into one image of the sequencer; (4) the large size of the datasets, containing millions of points; and (5) inconsistencies between the datasets in the form of missing points and positional errors.

Therefore, even though alignment of sequencing and fluorescence datasets has been done before (Buenrostro et al. 2014; Jung et al. 2017; She et al. 2017), it proved challenging to extend the method to single-molecule fluorescence imaging. Hence, we set out to make a procedure to perform the alignment even under the aforementioned challenging conditions. Here we show that we can accurately register the point sets obtained from single-molecule fluorescence and sequencing experiments using a three-step procedure that progresses from global to local registration. After registration, point correspondence is determined by setting a distance threshold based a mathematical model fit to the registered data. Our approach will not only be useful for labs combining sequencing with fluorescence assays—either at the single-molecule level or at the cluster level—but can be used for combining data from any other imaging methods.

## RESULTS

To register single-molecule data with sequencing data, we employed a three-step approach (Figure 1). First, since the imaging parameters of the sequencer were unknown, we needed to find the global rotation and scaling parameters, specific to the fluorescence microscope and sequencer. Because of the large search space for these variables, this registration had to be performed on a low-density dataset. Second, once the rotation and scaling parameters were known, a fast, coarse registration was performed for full datasets by determining the translation of each sequencing image, called a tile, within the single-molecule data. Third, as each fluorescence field of view may have small deviations with respect to the course registration, the registration for each individual field of view was fine-tuned, allowing small variations in translation, rotation and scaling. After registration, single-molecule to sequence correspondence was determined using a distance threshold, while at the same time requiring unique matches.

**Figure 1:**
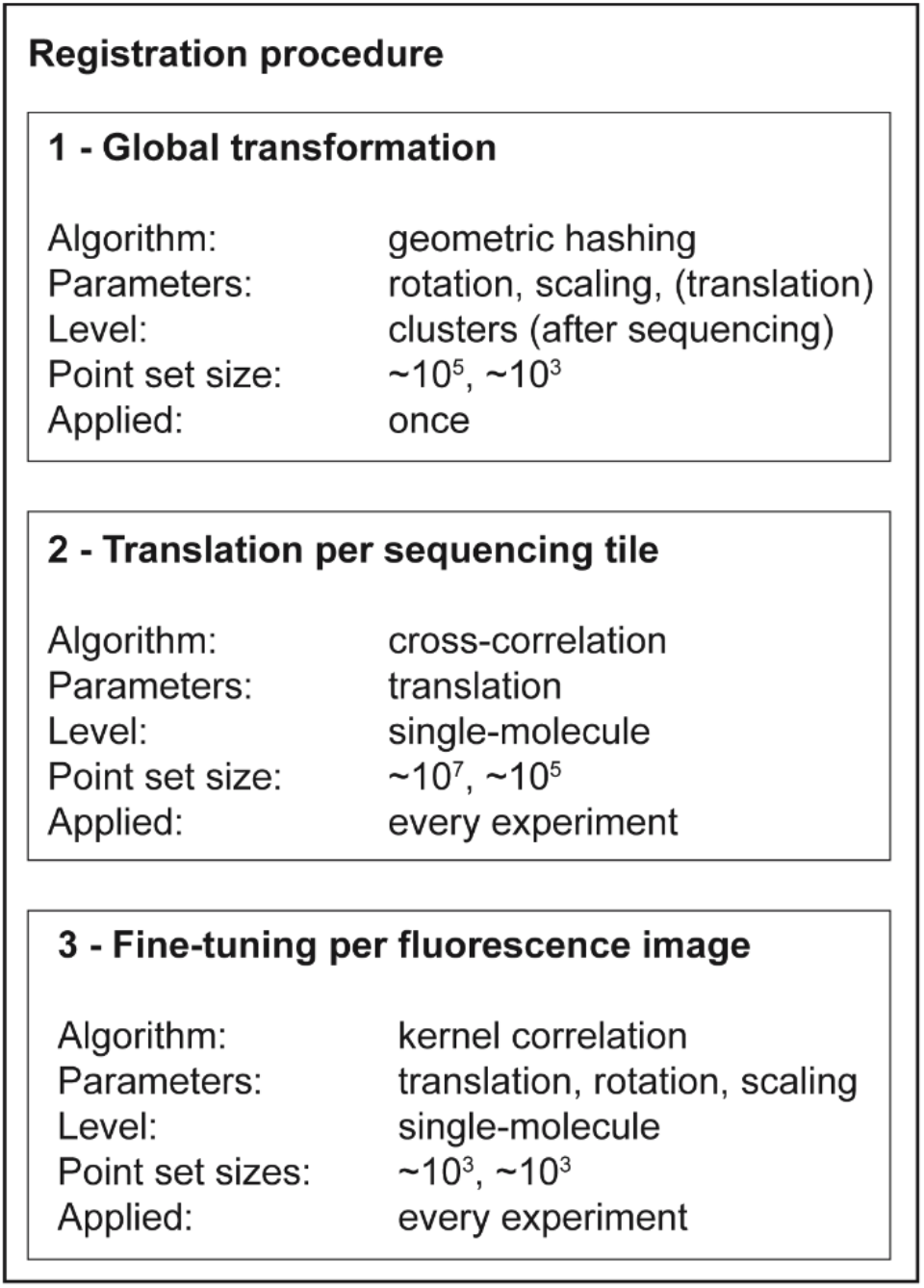
Overview of the three-step registration procedure.

### Global transformation

The global transformation between the fluorescence microscopy images and the sequencer output is determined using a dataset with fiducial markers after sequencing on the cluster level. To this end, we employed an adapted version of the geometric hashing algorithm (Wolfson and Rigoutsos 1997; Lang et al. 2010), which has been applied to find the location of star constellations among all the stars in the night sky (Lang et al. 2010). Unlike other point set registration algorithms (Zhu et al. 2019; Gang Wang 2024), geometric hashing is well suited for a large difference in dataset size, i.e. when finding a smaller point set in a larger point set (Wolfson and Rigoutsos 1997; Lang et al. 2010). Furthermore, it can handle positional noise and missing points (Wolfson and Rigoutsos 1997; Lang et al. 2010).

Our adaptation of the algorithm compares the shapes of point quadruplets, selected from the point sets and finds the common transformation of alike quadruplets (Figure 2A). After finding quadruplets within a point set, two of the four points in each quadruplet are used for transforming the quadruplet to a transformation-invariant coordinate system. The coordinates of the remaining two points in each quadruplet are then used to construct a hash table, capturing the shape of each quadruplet in four values that are stored in a hash table row. By comparing the hash tables of the to-be-matched point sets, the entries with resembling shapes are determined. When examining the transformation parameters that belong to pairs of resembling quadruplets, the correctly matched quadruplet pairs will cluster, as they all have similar transformations; on the other hand, the incorrectly matched pairs will be spread randomly across a larger space.

**Figure 2:**
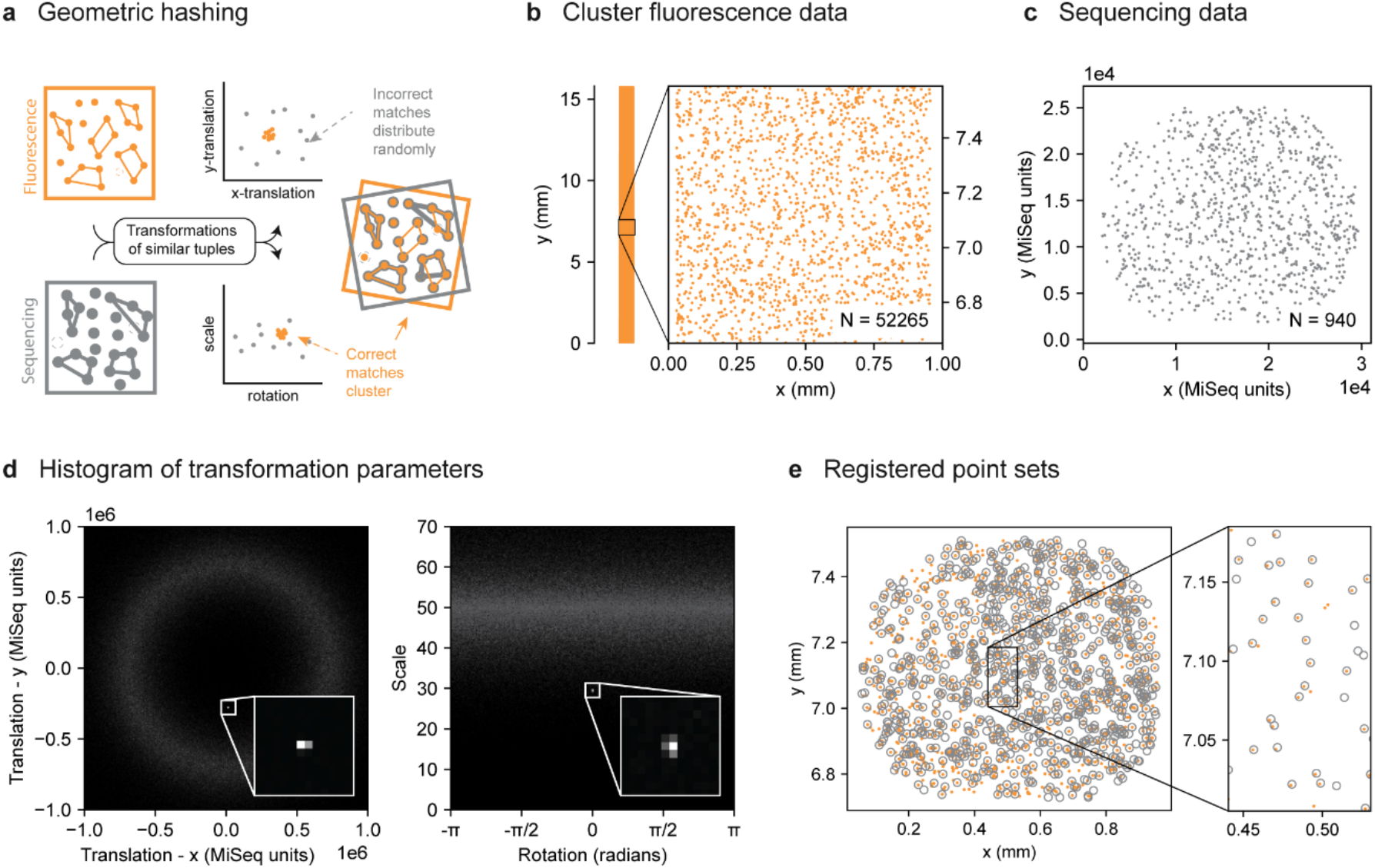
Global transformation between fluorescence microscope and sequencer coordinate systems. (A) Schematic of the geometric hashing algorithm. From the single-molecule fluorescence and sequencing point sets quadruplets of points are selected. For similarly-shaped quadruplets, the transformation parameters are calculated for the transformation between the single-molecule fluorescence and sequencing datasets. Correctly matched quadruplets all have similar transformations and thus cluster together; on the other hand, incorrectly matched quadruplets have more randomly distributed transformations. By finding the cluster, the correct transformation is obtained. (B) Fluorescence dataset obtained after sequencing, consisting of the DNA cluster locations with a specific marker sequence. Locations were extracted from the ∼7000 images obtained by scanning an area of ∼1×16 mm^2^. The coordinates were stitched together based on the automated stage coordinates. “N” indicates the full dataset size. (C) Sequencing dataset obtained from the coordinates of the marker sequence on a single sequencing tile. The coordinates were used as reported in the fastq file. The area of a single tile is ∼0.8×1 mm^2^.(D) Two-dimensional histograms of transformation parameters - translation (left) and rotation and scale (right) - found for similarly-shaped quadruplets in both datasets. (E) Result of the point set registration using geometric hashing. Orange points represent the cluster fluorescence data, grey open circles indicate the transformed sequencing data.

Although the algorithm can be used to only find the scaling and rotation parameters, the use of additional parameters will result in a clearer cluster, as the incorrectly matched pairs will be distributed over more dimensions. Therefore, it is beneficial to also add translation, which, for application to the single-molecule and sequencing data, means that only one tile can be matched at a time.

While geometric hashing can handle large differences in dataset size, the algorithm is limited by the hash table size and thus by the number of points used as input. In addition, higher percentages of missing points or larger positional errors would require larger hash tables, as fewer correct quadruplets can be matched between the points sets. To reduce the point set size and the number of missing points, this registration step was performed using DNA sample with a low concentration of a fiducial marker that was imaged after sequencing, at which point the single molecules were amplified into high-intensity clusters. This is possible since in the current step we are determining the global transformation between the data obtained on the specific sequencing instrument and the data obtained on the fluorescence microscope, which should be applicable to all experiments using this combination.

To obtain the dataset of fiducial markers, we sequenced a DNA sample that could be recognized in both the sequencing data and the fluorescence images, taken after sequencing upon hybridizing a fluorescent probe. In this way, the number of points was reduced approximately 250 fold with respect to a full dataset (from ∼250,000 to ∼1,000 sequences per tile), which was essential for finding a correct registration.

Alternatively, the size of the fluorescence point set can be reduced by using of a smaller sequencing chip, e.g. the MiSeq Nano, covering an area of roughly 2 mm^2^ instead of 16 mm^2^. Using the fiducial markers, the fluorescence and sequencing datasets of the entire chip, shown in Figure 2B and C, were registered in several minutes on a desktop computer. The distribution of transformation parameters for similarly shaped quadruplets showed a clear cluster for translation, rotation and scaling (Figure 2D). The resulting registration (Figure 2E) establishes the relation between the coordinates of the sequencer and the pixels in the fluorescence microscope, which was used as a starting point for subsequent experiments performed using these devices.

### Translation per tile

When the rotation and scaling for the specific combination of sequencer and microscopy setup are determined in **step 1**, the faster cross-correlation can be applied in subsequent experiments to find the location (translation) of each sequencing tile in the stitched single-molecule data. As cross correlation is calculated for images and not point sets, the locations exported by the sequencer and the coordinates found in the single-molecule images are converted into synthetic images (Figure 3A and B), where the transformation found in **step 1** is already applied. In the images, each point is represented by a two-dimensional Gaussian. For reconstructing an image of the sequencing data, only the sequences with a fluorescent label are used, since these are the only ones visible in the single-molecule images. While it is possible to obtain the raw images from the sequencer for alignment, in practice their use will be more cumbersome as the images would require resizing and would take up considerable disk space. Furthermore, cluster shape in the raw images will be different from the point spread function of single molecules.

**Figure 3:**
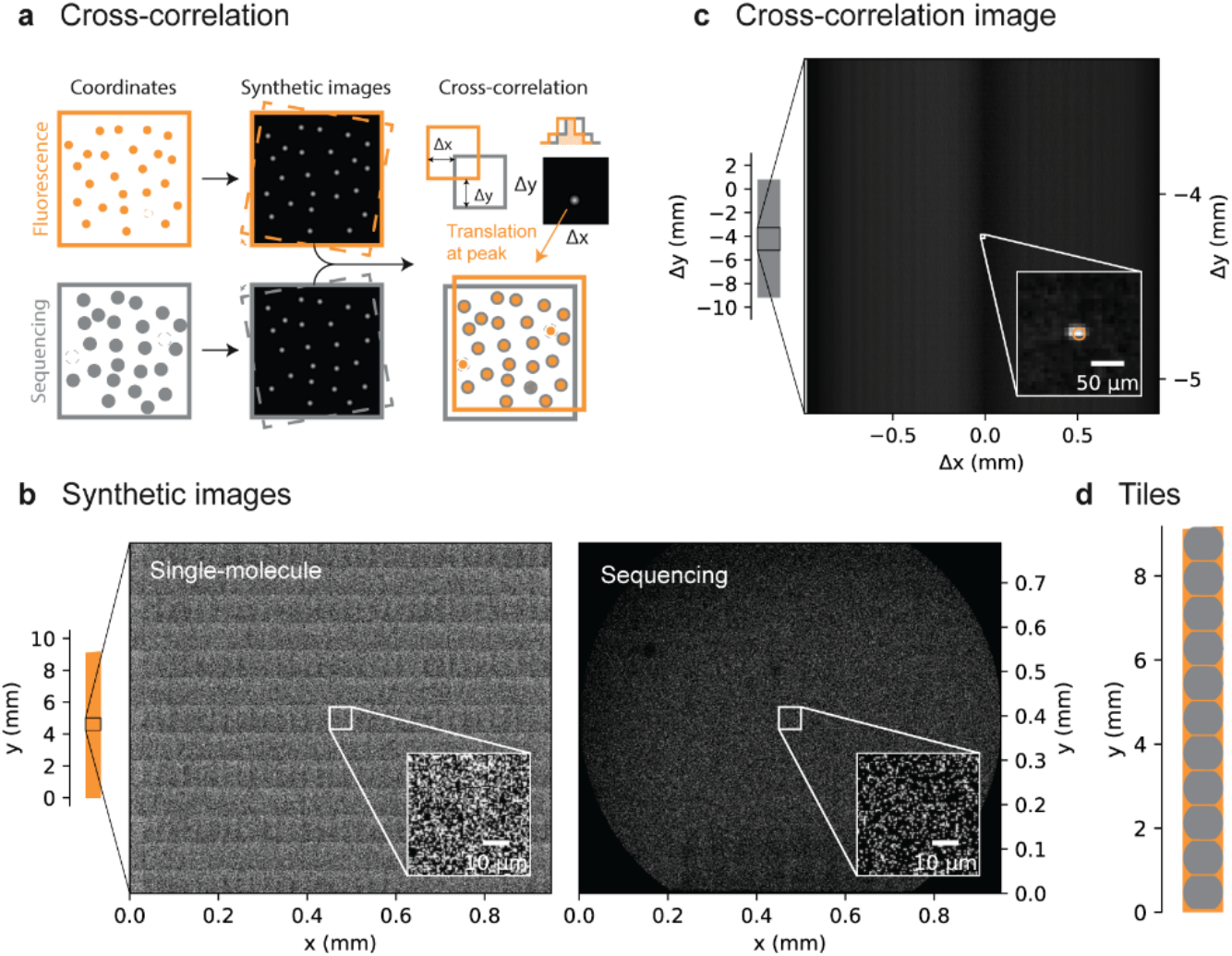
Translation per sequencing tile. (A) Schematic of the algorithm to cross-correlate the datasets. The point sets are first transformed using the scale and rotation found for the specific fluorescence and sequencing setups. The point sets are subsequently converted to images using two-dimensional Gaussians at the point coordinates. The images are then cross-correlated, after which the peak in the cross-correlation plot indicates the correct translation. (B) Synthetic images of the single-molecule dataset (left) and the sequencing dataset (right). The single-molecule data consists of all single-molecule locations from ∼4700 images obtained from scanning an area of ∼1×9 mm. The sequencing data consists of all sequences with a fluorescent label from a single tile. (C) Cross-correlation image after background subtraction. Inset shows the correlation peak. (D) Overview of the registration of the 11 sequencing tiles (grey) to the single-molecule fluorescence dataset (orange).

The cross-correlation image of the two synthetic images (Figure 3C) shows a sharp peak indicating the correct translation parameter. To verify the result, the same correlation was performed with one of the datasets mirrored, which did not show such a clear peak. All 11 neighboring tiles were found in the single fluorescence dataset with approximately equal spacing in the y-direction and similar x-position (Figure 3D). Thus, using the cross-correlation peak and the positions of multiple tiles, correctly matching tile locations could clearly be distinguished from random matches. Thereby, cross correlation enables fast registration of datasets that are too large to be processed with geometric hashing in **step 1**.

### Fine-tuning per fluorescence image

After registration to the sequencing tile, additional registration is necessary as the image resolution and Gaussian width used for creating the synthetic images in **step 2** limit the accuracy of the obtained translation parameters. Moreover, transformations may also vary slightly between individual fields of view (FOVs), due to positional variation by slight movements of the flow cell in the holder during scanning; or due to variations in scaling as a result of aberrations in the larger sequencing image. Therefore, additional fine-tuning is performed on each fluorescence image separately using a kernel correlation algorithm that allows slight variations in translation, rotation and scaling (Figure 4A).

**Figure 4:**
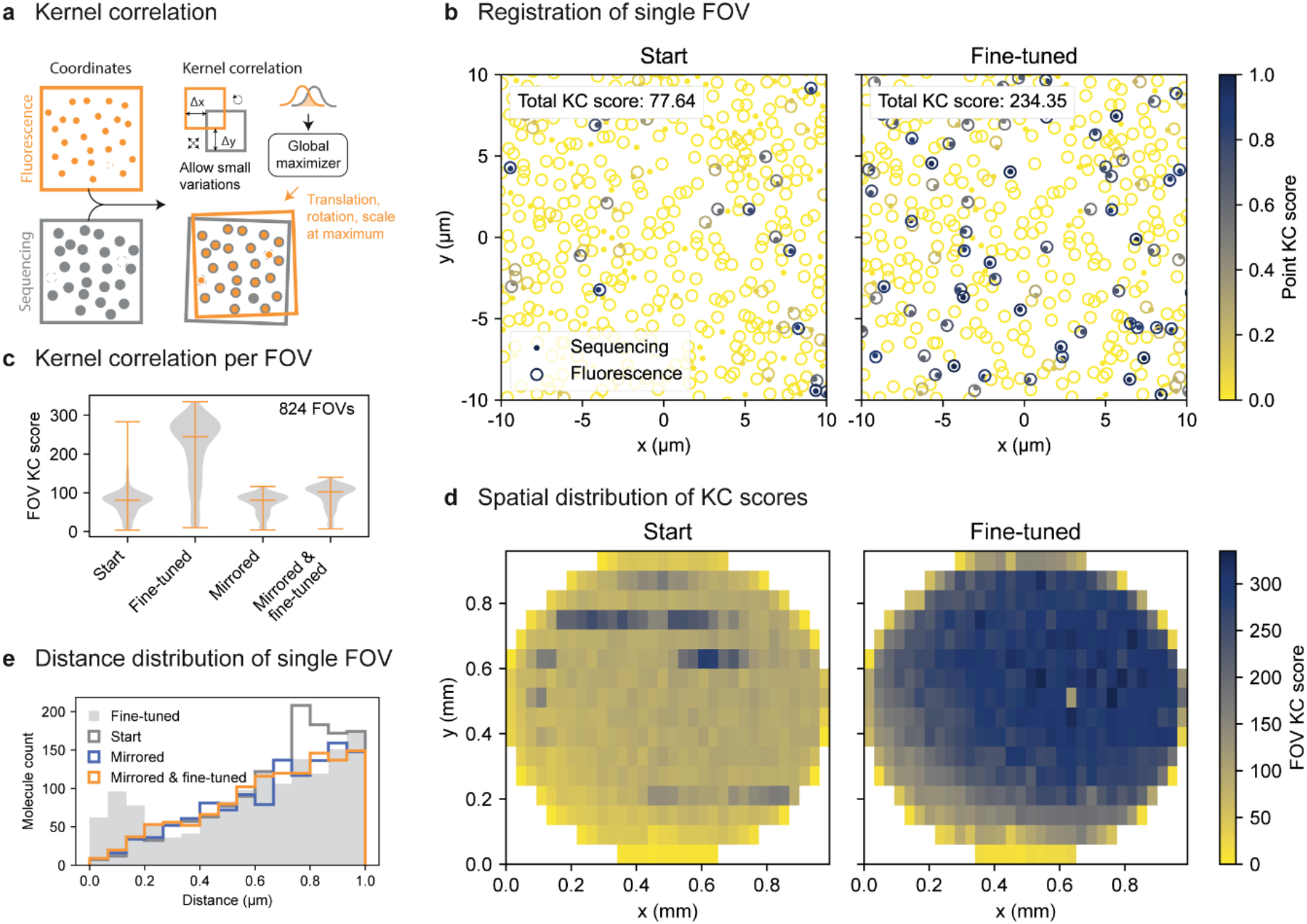
Fine-tuning per fluorescence image. (A) Schematic of the kernel correlation algorithm. For a specific transformation the kernel correlation between two points is calculated from the overlapping volumes of the Gaussians at their respective positions. By allowing small variations in translation, rotation and scale, for the transformation of each individual fluorescence field of view (FOV) the total kernel correlation for FOV is maximized to find the correct transformation. (B) Kernel correlation for a single FOV. Open circles indicate transformed single-molecule data, dots indicate sequencing data. The color scale for each individual point indicates the sum of the kernel correlation scores of that point to all other points in the other dataset; blue indicates a high score. (C) The distribution of kernel correlation scores. Each data point is the summed score for all point pairs in a FOV. The outer horizontal bars indicate the data extremes, the inner bar shows the median value. (D) Spatial distribution of total kernel correlation scores within a sequencing tile. Each rectangle represents a single FOV. (E) Histogram of the pairwise inter-point set distances for a single FOV. “Start” indicates the situation right after cross-correlation in **step 2**. For the “fine-tuned” dataset the transformations were adjusted to maximize the kernel correlation score. The “mirrored” dataset was reflected with respect to the “fine-tuned” case and thus functions as a negative control for registration. For the “mirrored & fine-tuned” data the reflected dataset was additionally fine-tuned using the same settings used for the “fine-tuned” dataset.

The kernel correlation (KC) score for a combination of one fluorescence and one sequencing point was calculated from the overlap of Gaussians centered at their positions (a mathematical description can be found in the Supplementary material ). The total score for a FOV was calculated as the sum of scores obtained from all combinations of fluorescence and sequencing points. The low KC scores for the transformation obtained from step 2, shows that the majority of FOVs was not registered properly (Figure 4B, C and D - start). For comparison with a dataset containing no correct match, scores were calculated for the case where one of the point sets was mirrored (Figure 4C - mirrored). This shows that at the start of step 3 - before fine-tuning - the majority of the FOVs have a KC score comparable with unmatched datasets.

Fine-tuning the transformation using kernel correlation dramatically improved the registration, as shown by the increased kernel correlation scores (Figure 4B, C and D - fine-tuned). Applying the same procedure to the mirrored dataset only slightly increases the KC score, thus indicating accurate registration for the regular dataset.

The validity of the registration can be observed in the pairwise inter-point setpoint distance distribution for a single FOV (Figure 4E). The “fine-tuned” registration displays a clear peak for short distances in the pairwise distance distribution, indicating accurate registration. A similar peak, although at a larger distance, can be seen before fine-tuning, showing that the initial alignment for this FOV deviated by approximately 0.8 µm. On the other hand, the “mirrored” and “mirrored & fine-tuned” cases (representing two uncorrelated point sets) show a nearly linear increase in the distribution, without peaks.

### Point correspondence

After obtaining an accurate registration, we determined point correspondence, identifying for each single-molecule fluorescence point to which sequencing point (if any) it corresponds. To do this we used a fixed threshold, estimated from the parameters obtained by fitting a model to the distribution of measured point distances. The model (Figure 5A) assumes a complete point set of size *N* from which the single-molecule and sequencing point sets are sampled with efficiencies *α*_*sm*_ and *α*_*seq*_, giving rise to point sets with sizes N_sm_ and N_seq_. Of these α_sm_N_seq_ = α_seq_N_sm_ = α_sm_α_seq_N = N_corr_ points occur in both datasets. In addition, it is assumed that the sampled point sets have a normally distributed positional error 𝒩 (σ^2^) with respect to the original point set. Such error can originate, for example, from a physical change of molecule position between obtaining the single-molecule and sequencing datasets, from noise or aberrations in the measurement, and from inaccuracies in localization during analysis. Lower positional error and higher sampling efficiencies will result in easier registration and more accurate correspondence.

**Figure 5:**
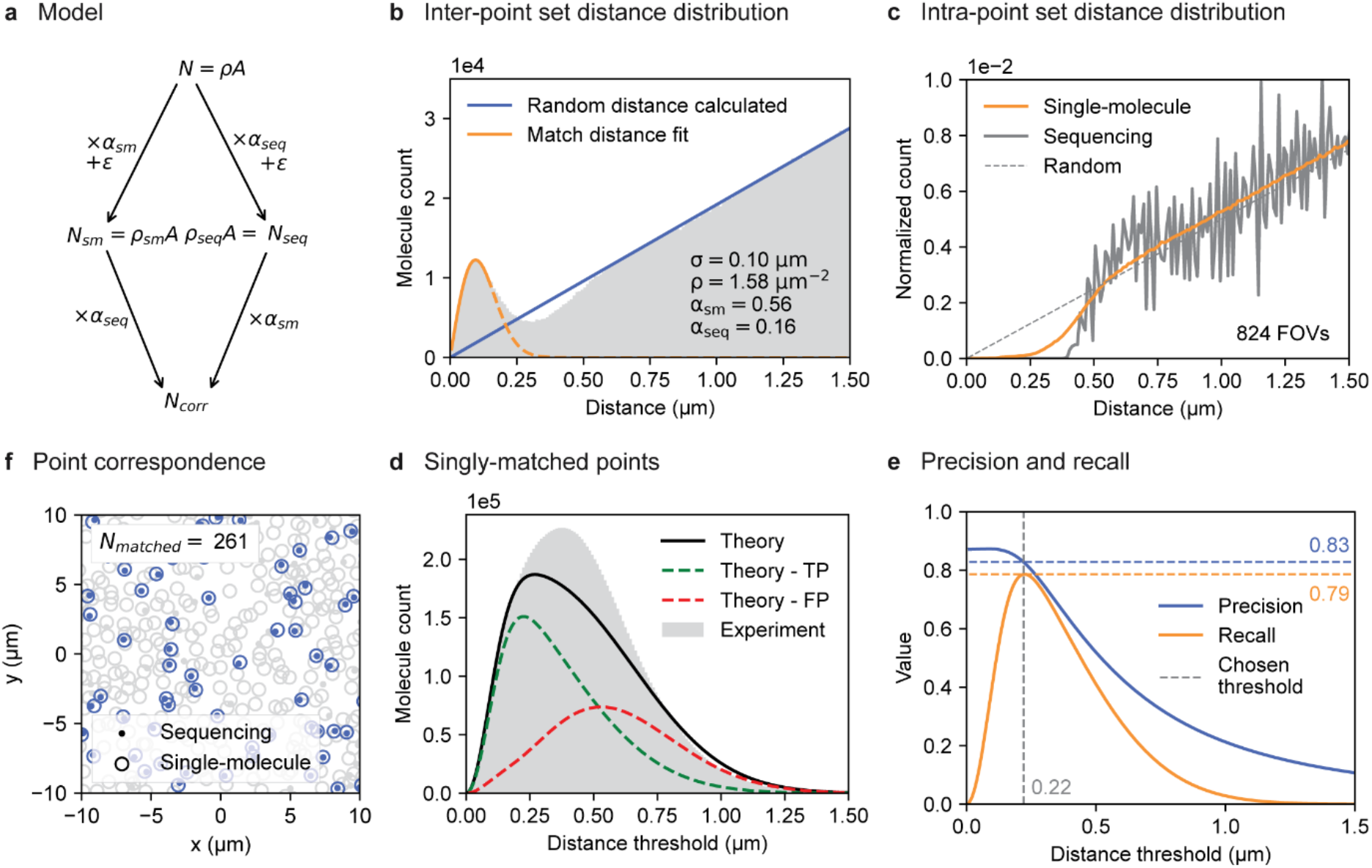
Point correspondence. (A) Schematic of the model used for estimating parameters of the point set. Starting out with all N points, a fraction α_sm_ will be visible in the single-molecule data and a fraction α_*seq*_ in the sequencing data, resulting in N_sm_ and N_seq_ points in these datasets respectively. The number of points can also be written as the density ρ times the imaged surface area A. A fraction α_seq_ of N_sm_ will be matched to a fraction α_sm_ of N_seq_, resulting in N_corr_ matched points. (B) Distribution of distances between the points of the single-molecule and sequencing datasets. The orange line indicates the fit of the distance distribution for corresponding points; only the solid section is actually fit. The blue line indicates the contribution of the randomly occurring point pairs. Data originate from 824 fields of view (FOVs). (C) Distribution of the distances between points within the single-molecule point set (orange) and the sequencing point set (grey). The dotted line indicates a random distribution, which is proportional to the circumference of a circle and thus to the distance *r*. Data originate from 824 fields of view (FOVs). (D) The number of singly-matched point pairs for a specific distance threshold, i.e. a single-molecule point is matched to one and only one sequencing point and vice versa. The grey histogram shows the values determined from experiments. The solid black line indicates the theoretical prediction based on the fit parameters found in (B). Green and red dashed lines indicate the theoretical contributions of true positive (TP) and false positive (FP) point matches. (E) Precision (blue) and recall (orange) as determined from the theoretical true positive and false positive contributions in (D). For the chosen threshold (grey dashed line) values for precision and recall (blue and orange dashed lines respectively) are indicated. (F) Point correspondence determined for a single field of view (FOV). Open circles indicate the transformed single-molecule data, the dots indicate sequencing data. Grey points are unmatched, blue points are matched. N_matched_ indicates the number of matched points.

The model yields a Gaussian distribution for distances between corresponding points and a linearly increasing distribution for distances between randomly paired points (See Methods and Supplementary material). The obtained distance histogram (Figure 5B) indeed shows these two populations: a Gaussian peak for smaller values of r going over in a linear relationship for larger values of r. Fitting the left side of the histogram with the distance distribution for corresponding points using σ and N_corr_ as free parameters (Figure 5B) yield a good agreement with the model. In addition, the right side of the histogram corresponds well with a random distribution based on the measured densities ρ_sm_ and ρ_seq_ of the two point sets. However, for r around 0.4 µm the measured values are lower than expected. This deviation is likely caused by the point detection algorithms in both single-molecule and sequencing analysis, which remove points that are located closely together or possibly combine them into a single point. This can be seen in the distribution of distances within each point set (Figure 5C), where the number of point pairs with distances smaller than approximately 0.5 µm is lower than expected based on a fully random distribution. In turn, this affects the inter-point set distance distribution, because detection of neighboring single-molecule and sequencing points makes it unlikely to find additional points close by. Although this effect is easy to detect, it is hard to theoretically quantify. Therefore, we will use the fitted distribution as an upper boundary for the error.

Using the parameters estimated from the distance distribution (Figure 5B) the matching error could be calculated and an appropriate distance threshold could be chosen. All model parameters were determined, either directly from the experimental data (N_sm_, N_seq_ and A) , from fitting the distance distributions(σ and N_corr,_) or from derivations of other parameters (α_sm,_ α_seq_, ρ_sm_, ρ_seq_, ρ and N). We additionally imposed that the points were singly matched, i.e. a single-molecule fluorescence point should have precisely one match in the sequencing data and vice versa. The theoretically expected and experimentally determined numbers of singly-matched points for different distance thresholds are shown in Figure 5D. The plot shows a rise for smaller thresholds as the tolerance for positional offsets increases, and a fall to zero for larger distance thresholds, where the probability to detect multiple points within the threshold increases. For small and large thresholds the theoretical curve follows the experimental values closely; however, for intermediate thresholds the theory underestimates the experiment. Again, the cause could be the absence of small point distances within a point set (Figure 5C). Using the theoretical values of true and false positive probabilities the precision and recall were calculated (Figure 5E). Finally, the distance threshold with the largest recall – 0.22 µm – was chosen giving a precision of 83% and recall of 79%. This resulted on average in 238 corresponding points per field of view, out of a total of 1473 single-molecule fluorescence points and 465 sequencing points. An example is shown in Figure 5F.

## DISCUSSION

Here we provided a methodical approach for point set registration in experiments combining fluorescence and sequencing data. We have shown that single-molecule fluorescence and sequencing data can be accurately registered using a three-step approach that progresses from global to local registration: (1) finding the global transformation for the specific microscope using geometric hashing, (2) determining the translation of each sequencing tile using cross-correlation, and (3) fine-tuning the transformation of each single-molecule fluorescence image using kernel correlation. After these steps, point correspondence is established by setting an absolute distance threshold and ensuring that there is only one possible match within the threshold.

While registration of fluorescence microscopy and sequencing data has been done previously (Buenrostro et al. 2014; Jung et al. 2017; She et al. 2017), it remained a challenge to extend it to single-molecule fluorescence data. Particularly, the large number of missing points in the single-molecule data obtained before sequencing increased the difficulty of registration compared to registration of cluster-level data obtained after sequencing. Nevertheless, the kernel correlation algorithm could successfully provide proper registration even under these challenging conditions.

To make this approach readily accessible, we constructed a python library named MatchPoint for point set registration, containing, among others, the algorithms used here—geometric hashing, cross-correlation and kernel correlation—and additional functions for obtaining match statistics and plots. Although in the current work we focus on the combination of fluorescence and sequencing data, we envision that a similar approach can be more generally applied to perform point set registration for combinations of other microscopy systems. By providing accessible tools for point set registration, we aim to facilitate the implementation of combined microscopy methods across various scientific applications.

## EXPERIMENTAL PROCEDURES

### Dataset with fiducial markers at the cluster level

The dataset with fiducial markers on the cluster level was obtained as described in (Severins et al. in review). In short, a specific DNA sequence called the mapping sequence was inserted at low concentration into a DNA library and subsequently sequenced on an Illumina MiSeq machine. After sequencing the secondary DNA strand, produced during the sequencing-by-synthesis process, was removed. Then, a fluorescently-labelled DNA probe complementary to the unique part of the mapping sequence was hybridized to the cluster DNA. The surface of the MiSeq flow cell was then scanned using an objective-type total internal reflection fluorescence microscope with a field of view size of 64 by 32 µm. From the images the positions of high intensity clusters, that reported the presence of the mapping sequence, were extracted. The found coordinates in the various images were stitched together based on the coordinates reported by the automated stage, which resulted in the fluorescence point set of fiducial markers. To obtain the positions of the mapping sequence in the sequencing data, the sequences were aligned to the reference sequences corresponding to the samples present in the DNA library. Using the alignment, the cluster positions specific to the mapping sequence were extracted. As the sequencer can image both top and bottom surfaces of the flow cell, only sequencing data was included from the side that was imaged using the fluorescence microscope. The result was the sequencing point set of the fiducial marker.

### Full single-molecule and sequencing dataset

The full single-molecule fluorescence and sequencing datasets were obtained as described in (Severins et al. in review). In short, a library of fluorescently-labelled DNA was immobilized on a MiSeq flow cell. The surface of the flow cell was scanned using an objective-type total internal reflection fluorescence microscope with a field of view size of 64 by 32 µm. From the images the locations showing single-molecule fluorescence were determined. The found coordinates in the various images were stitched together based on the coordinates reported by the automated stage, the result was the single-molecule fluorescence point set. Subsequently, the flow cell was sequenced on an Illumina MiSeq sequencer and the acquired data was aligned to the reference sequences of the samples in the library. The sequencing point set was then obtained by selecting the cluster locations of the fluorescently-labelled sequences from the side of the flow cell that was imaged on the fluorescence microscope.

### Geometric hashing

Our implementation of the geometric hashing algorithm is an adaptation of previously published methods (Wolfson and Rigoutsos 1997; Lang et al. 2010). The aim of the geometric hashing algorithm was to find the transformation *T* that correctly registered the source point set P^S^ = {p_1_^S^, …, p^S^_NS_} and the destination point set P^D^ = {p_1_^D^, …, p^S^_ND_}. To this end, quadruplets were selected from each point set Q_i_ = {p_j_, p_k_, p_l_, p_m_} where j ≠ k ≠ l ≠ m. This was done by first finding all pairs of (outer) points p_j_ and p_k_ with a distance smaller than the set threshold:

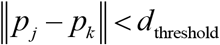

Then (inner) points p_l_ and p_m_ were chosen from the points within the circle that passed through p_j_ and p_k_, i.e.

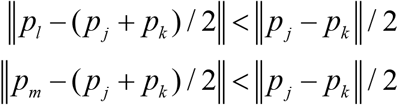

where (p_j_ + p_k_)/2 is the center of the circle and (p_j_ - p_k_)/2 is the radius. Quadruplets were made for all combinations of points within the circle.

To construct a transformation-invariant hash table with entries describing the quadruplet shape, the quadruplets were transformed to an invariant coordinate system using the two outer points as a basis (Lang et al. 2010). This was done by placing the outer points *p*_*j*_ and *p*_*k*_ on the coordinates (0,0) and (1,1) using transformation T_i_, i.e. (0,0) = T_i_ p_j_ and (1,1) = T_i_ p_k_. Since two basis points were used, only translation, rotation and scaling can be determined, effectively making T_i_ a similarity transformation (excluding reflection). The hash table values h_i_ were then obtained by transforming the remaining, inner points of the quadruplet, i.e. p′_l_ = T_i_ p_l_ and p′_m_ = T_i_ p_m_, and obtaining their x and y coordinates: h_i_ = (p′_l,x_ , p′_l,y_, p′_m,x_, p′_m,y_). As it is still possible to form different hash table values from the same quadruplet by switching j and k, or l and m, the following additional criteria were added to establish a unique point order:

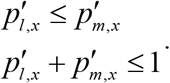

Hash tables (H^S^ and H^D^) accompanied by the tables containing the transformation matrix of each entry (T^S^ and T^D^) were constructed for both the source and destination point sets. Hash table entries in the two datasets were compared and entries with distances smaller than a threshold were selected,

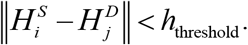

Subsequently, the transformation matrices describing the transformation from source to destination were calculated for each of the paired hash table entries:

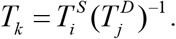

Then translation, rotation and scaling (t_x_, t_y_, r and s) parameters were extracted from the transformation matrices. For correctly matched hash table entries the transformation parameters are similar and thus cluster together, while for incorrect matches the parameters will be (partially) random and are thus spread out. The cluster of similar transformations was found by normalizing the values each of the four parameters separately, e.g.

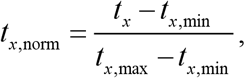

to give each parameter similar weight, and by then finding the point(s) with the highest number of local neighbors within a threshold. The final transformation was determined by taking the average of the transformation parameters over the found points.

The geometric hashing algorithm was implemented in the Python programming language using NumPy, SciPy and scikit-image libraries. For efficient lookup, point sets and hash tables were stored in kd-tree data structures.

### Cross-correlation

Fast Fourier transform based algorithms for cross-correlation are efficient for registering translated images. Therefore, the coordinates in each point set were converted to a synthetic image. This was done by first applying the scaling and rotation parameters found in step 1 to the point-set coordinates. Subsequently, the transformed coordinates were converted to pixel indices by scaling the coordinate values to the desired pixel range (0 to N_pixel,max_) and by rounding to integers:

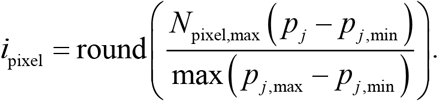

Then a Boolean matrix B was constructed where the values at the found pixel indices were set to 1 and the remaining indices were set to 0. The final synthetic image I was then constructed by convolution of a Gaussian kernel

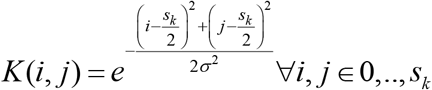

(with kernel size s_k_ =7 pixels and σ = 1 pixel) with the Boolean matrix:

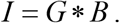

To find the location of the source image into the destination image, cross correlation was performed:

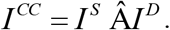

To reduce the variation in background intensity, a minimum filter was applied to the cross-correlation image:

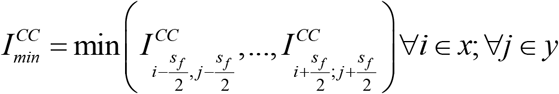

(where filter size s_f_ = 2 s_k_ was used) and the result was subtracted:

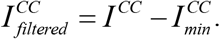

Then the pixel with the highest correlation value was found and its index was converted back to a translation value. The found value was added to the original transformation from step 1.

Convolution and cross-correlation were performed using the SciPy signal package.

### Kernel correlation

The kernel correlation (Tsin and Kanade 2004) between two points p_i_ and p_j_ was defined as

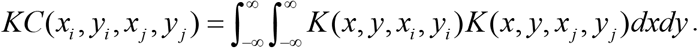

In our case a Gaussian kernel

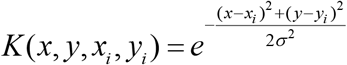

was used. The Kernel correlation then became a function of the distance d_i,j_ between the two points:

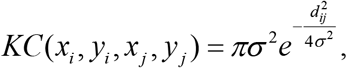

where

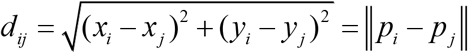

(for a derivation, see the Supplementary material).

The total kernel correlation value for the two point sets is the sum for all point combinations:

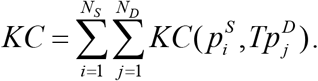

To calculate the total kernel correlation value in practice, i.e. with sufficient speed, the point sets were stored in kd-tree data structures and distances between the points in the two point sets were only determined if they were closer than a threshold:

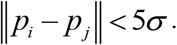

The total kernel correlation value was maximized using the differential evolution algorithm (Storn and Price 1997), a global optimizer implemented in the SciPy library.

### Point correspondence

Point set correspondence was determined by setting a threshold based on a statistical model for randomly distributed point sets. The statistical model assumes a “perfect” point set of N points from which the single-molecule fluorescence and sequencing point sets were sampled with efficiencies α_sm_ and α_seq_ to give N_sm_ and N_seq_ points, respectively. At the same time this means that the number of corresponding points between the single-molecule and sequencing point sets is

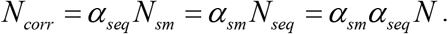

In addition, the sampled point sets were assumed to have normally distributed positional errors 𝒩 (σ_sm_^2^) and 𝒩 (σ_seq_^2^) with respect to the original point set.

For corresponding points from the single-molecule and sequencing point sets which are both subject to their own error, this leads to a distribution d_corr_ over distance r:

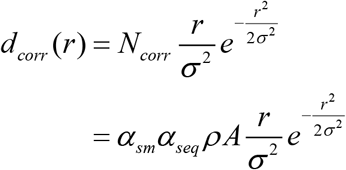

where ρ is the density of the perfect point set, A is the area imaged and σ^2^ = σ_sm_^2^ + σ_seq_^2^, is the combined variance of the single-molecule and sequencing datasets. On the other hand, the distance distribution for randomly matched points is proportional to the circumference of a circle:

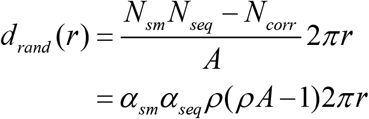

The binned versions of the these distance distributions with binwidth Δ*r* are

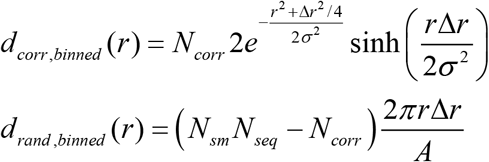

While the values of N_sm_, N_seq_ and A could be directly obtained from the point sets, the values of N_corr_ and σ were determined by fitting the distribution for corresponding points d_corr_ to the Gaussian part of the measured distance histogram. Here we did not use the sum of the distributions for corresponding and random points for fitting as it did not describe the data well for smaller distances. There, the theoretical number of random matches is likely overestimated due to the removal of nearby points during extraction of the point coordinates from the sequencing and single-molecule images. From the fitted parameters, in turn, α_sm_, α_seq_, ρ_sm_, ρ_seq_, ρ and N were calculated.

To determine point correspondence, a distance threshold was set and it was additionally imposed that the points are singly matched, i.e. a single-molecule point should have one and only one match in the sequencing data and vice versa. The number of singly-matched points found for a specific distance threshold can be measured and simultaneously estimated from theory using the probability for finding corresponding and random points within a distance threshold:

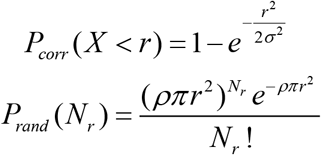

A derivation of the distributions can be found in the Supplementary material. Using these relations the probability of true positives P_TP_, false positives P_FP_, false negatives P_FN_ and true negatives P_TN_ can be determined. Where a positive was defined as the case where both of the corresponding single-molecule fluorescence and sequencing points were present, while negative was defined as the case where either or both of the points were absent. For example, a true positive meant that both corresponding single-molecule and sequencing points were present, that they were closer than the distance threshold *R* and, additionally, that no other points were located within the threshold:

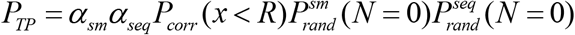

Similar, but more complex relations were derived for P_FP_, P_FN_ and P_TN_, these can be found in the Supplementary material. From these theoretical values, the precision (fraction of identified point correspondences that is correct) and recall (fraction of actual point correspondences that is correctly identified) were determined

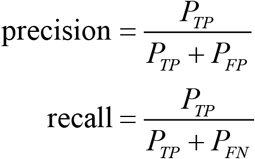

Using the calculated precision and recall for various distance thresholds an optimal distance threshold was chosen for point matching. Fitting was done using SciPy library.

## SUPPLEMENTAL INFORMATION DESCRIPTION

The supplemental information contains a derivation of the Gaussian kernel correlation function that is used in the implementation of the kernel correlation algorithm. Additionally, it contains derivations of distance distributions for random point sets and for corresponding point sets. These are then used to obtain the statistics of correspondence, i.e. true positive, false positive, true negative and false negative fractions, from which the precision and recall can be derived.

## ACKNOWLEDGEMENTS

The authors thank Carolien Bastiaanssen and Sung Hyun Kim for testing and feedback on the point set registration procedure.

## AUTHOR CONTRIBUTIONS

I.S. obtained and analyzed the single-molecule fluorescence and sequencing point set data. I.S. devised the adapted geometric hashing algorithm and developed the MatchPoint python package with implementations of geometric hashing, cross-correlation and kernel correlation algorithms. I.S. derived the correspondence statistics. J.v.N. and C.J. provided feedback throughout the project. I.S. wrote the manuscript with input from J.v.N. and C.J..

