## Supplementary material for "Point set registration for combining fluorescence microscopy methods"

**Derivation of the Gaussian kernel correlation function**

From the definition of the kernel correlation and using a Gaussian kernel:

$$
\begin{aligned}
KC(x_i, y_i, x_j, y_j) &= \int_{-\infty}^{\infty}\int_{-\infty}^{\infty} K(x, y, x_i, y_i)K(x, y, x_j, y_j)dxdy \\
&= \int_{-\infty}^{\infty}\int_{-\infty}^{\infty} e^{-\frac{(x-x_i)^2+(y-y_i)^2}{2\sigma^2}} e^{-\frac{(x-x_j)^2+(y-y_j)^2}{2\sigma^2}} dxdy \\
&= \int_{-\infty}^{\infty}\int_{-\infty}^{\infty} e^{-\frac{(x-x_i)^2+(y-y_i)^2+(x-x_j)^2+(y-y_j)^2}{2\sigma^2}} dxdy \\
&= \int_{-\infty}^{\infty} e^{-\frac{(x-x_i)^2+(x-x_j)^2}{2\sigma^2}} dx \int_{-\infty}^{\infty} e^{-\frac{(y-y_i)^2+(y-y_j)^2}{2\sigma^2}} dy
\end{aligned}
$$

We can write

$$
\begin{aligned}
&\int_{-\infty}^{\infty} e^{-\frac{(x-x_i)^2+(x-x_j)^2}{2\sigma^2}} dx \\
&= \int_{-\infty}^{\infty} e^{-\frac{x^2-2x_ix+x_i^2+x^2-2x_jx+x_j^2}{2\sigma^2}} dx \\
&= \int_{-\infty}^{\infty} e^{-\frac{2x^2-2(x_i+x_j)x+x_i^2+x_j^2}{2\sigma^2}} dx
\end{aligned}
$$

Then using:

$$
\int_{-\infty}^{\infty} e^{-ax^2+bx+c} dx = \sqrt{\frac{\pi}{a}} e^{\frac{b^2}{4a}+c}
$$

with

$$
\begin{aligned}
a &= 1/\sigma^2 \\
b &= (x_i + x_j)/\sigma^2 \\
c &= -(x_i^2 + x_j^2)/(2\sigma^2)
\end{aligned}
$$

we get

$$\begin{aligned}
\int_{-\infty}^{\infty} e^{-\frac{2x^2-2(x_i+x_j)x+x_i^2+x_j^2}{2\sigma^2}}dx &= \int_{-\infty}^{\infty} e^{-ax^2+bx+c}dx \\
&= \sqrt{\frac{\pi}{a}}e^{\frac{b^2}{4a}+c} \\
&= \sqrt{\frac{\pi}{1/\sigma^2}}e^{\frac{\left((x_i+x_j)/\sigma^2\right)^2}{4*1/\sigma^2}-\frac{x_i^2+x_j^2}{2\sigma^2}} \\
&= \sigma\sqrt{\pi}e^{\frac{(x_i+x_j)^2\sigma^2}{4\sigma^4}-\frac{x_i^2+x_j^2}{2\sigma^2}} \\
&= \sigma\sqrt{\pi}e^{\frac{(x_i+x_j)^2}{4\sigma^2}-\frac{2(x_i^2+x_j^2)}{4\sigma^2}} \\
&= \sigma\sqrt{\pi}e^{\frac{x_i^2+2x_ix_j+x_j^2}{4\sigma^2}-\frac{2(x_i^2+x_j^2)}{4\sigma^2}} \\
&= \sigma\sqrt{\pi}e^{\frac{-x_i^2+2x_ix_j-x_j^2}{4\sigma^2}} \\
&= \sigma\sqrt{\pi}e^{-\frac{(x_i-x_j)^2}{4\sigma^2}}
\end{aligned}$$

The kernel correlation value will then be

$$\begin{aligned}
KC(x_i, y_i, x_j, y_j) &= \int_{-\infty}^{\infty} e^{-\frac{(x-x_i)^2+(x-x_j)^2}{2\sigma^2}}dx\int_{-\infty}^{\infty} e^{-\frac{(y-y_i)^2+(y-y_j)^2}{2\sigma^2}}dy \\
&= \sigma\sqrt{\pi}e^{-\frac{(x_i-x_j)^2}{4\sigma^2}}\sigma\sqrt{\pi}e^{-\frac{(y_i-y_j)^2}{4\sigma^2}} \\
&= \pi\sigma^2 e^{-\frac{(x_i-x_j)^2+(y_i-y_j)^2}{4\sigma^2}} \\
&= \pi\sigma^2 e^{-\frac{r_{ij}^2}{4\sigma^2}}
\end{aligned}$$

where

$$r_{ij} = \sqrt{(x_i - x_j)^2 + (y_i - y_j)^2} = \left\| p_i - p_j \right\|$$

is the distance between the points.

**Derivation of distance distributions for random point sets**

*Probability density*

The probability density distribution for the location of a point randomly positioned within an area A is uniform:

$$f_{rand}(x,y) = 1/A$$

To determine the probability density for finding the point at a certain distance from a starting point we need to integrate over all probability densities at distance $r$ around the starting point:

$$\begin{aligned} f_{rand}(r) &= \int_{r^2=x^2+y^2} f_{rand}(x,y)dxdy \\ &= \int_{r^2=x^2+y^2} \frac{1}{A} rdrd\theta \\ &= \frac{1}{A} r \int_{-\pi}^{\pi} d\theta \\ &= \frac{2\pi r}{A} \end{aligned}$$

which is valid as long as $r$ does not pass the boundary of A.

*Cumulative probability distribution*

To find the cumulative probability function $F_{rand}$ to have the random point within radius r of the initial point, we have to integrate the probability density from 0 to r.

$$\begin{aligned} F_{rand}(r) &= \int_0^r f_{rand}(r')dr' \\ &= \int_0^r \frac{2\pi r'}{A} dr' \\ &= \frac{2\pi}{A} \int_0^r r'dr' \\ &= \frac{2\pi}{A} \left[ \frac{1}{2} r'^2 \right]_0^r \\ &= \frac{2\pi}{A} \left[ \frac{1}{2} r^2 - \frac{1}{2} 0^2 \right] \\ &= \frac{\pi r^2}{A} \end{aligned}$$

*Binned probability density*

To obtain the binned probability density for binwidth Δr we need to integrate over the bin:

$$
\begin{aligned}
f_{\text{rand, binned}}(r,\Delta r) &= \int_{r-\Delta r/2}^{r+\Delta r/2} f(r')dr' \\
&= \int_{0}^{r+\Delta r/2} f_{rand}(r')dr' - \int_{0}^{r-\Delta r/2} f_{rand}(r')dr' \\
&= F_{rand}(r+\Delta r/2) - F_{rand}(r-\Delta r/2) \\
&= \frac{\pi}{A}\left(r+\frac{\Delta r}{2}\right)^2 - \frac{\pi}{A}\left(r-\frac{\Delta r}{2}\right)^2 \\
&= \frac{\pi}{A}\left[\left(r^2+\frac{2r\Delta r}{2}+\frac{\Delta r^2}{4}\right)-\left(r^2-\frac{2r\Delta r}{2}+\frac{\Delta r^2}{4}\right)\right] \\
&= \frac{\pi}{A}\left[\frac{2r\Delta r}{2}+\frac{2r\Delta r}{2}\right] \\
&= \frac{2\pi r\Delta r}{A}
\end{aligned}
$$

*Probability*

To obtain the probability of finding a specific number of points within a circle we use the binomial distribution. Assuming there are N points within the area A,

$$
\begin{aligned}
P(N_r) &= \binom{N}{N_r}\left(F_{rand}(r)\right)^{N_r}\left(1-F_{rand}(r)\right)^{N-N_r} \\
&= \binom{N}{N_r}\left(\frac{\pi r^2}{A}\right)^{N_r}\left(1-\frac{\pi r^2}{A}\right)^{N-N_r}
\end{aligned}
$$

Using that for an infinite number of experiments the binomial distribution approaches the Poisson distribution if np = λ:

$$
\begin{aligned}
P(X=k) &= \binom{n}{k} p^k (1-p)^{n-k} \\
\lim_{n\to\infty} P(X=k) &= \frac{\lambda^k e^{-\lambda}}{k!}
\end{aligned}
$$

using

$$
\begin{aligned}
n &= N \\
k &= N_r \\
p &= \frac{\pi r^2}{A} \\
\lambda &= np = N\frac{\pi r^2}{A} = \rho\pi r^2
\end{aligned}
$$

where ρ is the density of the point set and λ is thus constant for a specific r, we get

$$
P(N_r) = \frac{(\rho\pi r^2)^{N_r} e^{-\rho\pi r^2}}{N_r!}
$$

For the $N_r = 0$ and $N_r = 1$ we thus get

$$
\begin{aligned}
P(N_r = 0) &= \frac{(\rho\pi r^2)^0 e^{-\rho\pi r^2}}{0!} = e^{-\rho\pi r^2} \\
P(N_r = 1) &= \frac{(\rho\pi r^2)^1 e^{-\rho\pi r^2}}{1!} = \rho\pi r^2 e^{-\rho\pi r^2}
\end{aligned}
$$

*Distance distribution*

To obtain the density of distances between the single-molecule and sequencing datasets, we need to multiply the probability density by the total number of point pairs that are randomly distributed. These are all the point combinations minus the combinations of corresponding points:

$$
\begin{aligned}
N_{rand} &= N_{sm}N_{seq} - N_{corr} \\
&= \alpha_{sm}\alpha_{seq}N(N-1) \\
&= \alpha_{sm}\alpha_{seq}\rho A(\rho A - 1)
\end{aligned}
$$

with $N_{sm}$ the number of single-molecule and $N_{seq}$ the number of sequencing points.

The distance distribution functions then become:

$$
\begin{aligned}
d_{rand}(r) &= \alpha_{sm}\alpha_{seq}N(N-1)\frac{2\pi r}{A} \\
d_{rand,binned}(r,\Delta r) &= \alpha_{sm}\alpha_{seq}N(N-1)\frac{2\pi r\Delta r}{A} \\
D_{rand}(r,\Delta r) &= \alpha_{sm}\alpha_{seq}N(N-1)\frac{\pi r^2}{A}
\end{aligned}
$$

**Derivation of distance distributions for corresponding point sets**

*Probability density*

We assume that the positions of single-molecule points and sequencing points are normally distributed in both x and y around their center at (0,0) due to positional errors, and that the standard deviations are $\sigma_x = \sigma_y = \sigma$. The 1D normally-distributed probability density function for one point then becomes:

$$f_{normal,1D}(x) = \frac{1}{\sigma\sqrt{2\pi}} e^{-\frac{x^2}{2\sigma^2}}$$

Finding the probability distribution for a specific pair of values $x_{sm}$ and $x_{seq}$ for a pair of single-molecule and sequencing points gives:

$$\begin{aligned} f_{corr,1D}(x_{sm}, x_{seq}) &= f_{normal,1D}(x_{sm}) f_{normal,1D}(x_{seq}) \\ &= \frac{1}{\sigma_{sm}\sqrt{2\pi}} e^{-\frac{x_{sm}^2}{2\sigma_{sm}^2}} \frac{1}{\sigma_{seq}\sqrt{2\pi}} e^{-\frac{x_{seq}^2}{2\sigma_{seq}^2}} \\ &= \frac{1}{2\pi\sigma_{sm}\sigma_{seq}} e^{-\left(\frac{x_{sm}^2}{2\sigma_{sm}^2} + \frac{x_{seq}^2}{2\sigma_{seq}^2}\right)} \\ &= \frac{1}{2\pi\sigma_{sm}\sigma_{seq}} e^{-\left(\frac{x_{sm}^2\sigma_{seq}^2}{2\sigma_{sm}^2\sigma_{seq}^2} + \frac{x_{seq}^2\sigma_{sm}^2}{2\sigma_{sm}^2\sigma_{seq}^2}\right)} \\ &= \frac{1}{2\pi\sigma_{sm}\sigma_{seq}} e^{-\frac{x_{sm}^2\sigma_{seq}^2 + x_{seq}^2\sigma_{sm}^2}{2\sigma_{sm}^2\sigma_{seq}^2}} \end{aligned}$$

With $d$ the distance between the points, then:

$$\begin{aligned} d &= x_{seq} - x_{sm} \\ x_{seq} &= d + x_{sm} \\ x_{seq}^2 &= (d + x_{sm})^2 \\ &= d^2 + 2dx_{sm} + x_{sm}^2 \end{aligned}$$

where we assume $x_{seq} > x_{sm}$. The alternate case ($x_{seq} < x_{sm}$) will be taken into account later.

Now we get

$$
\begin{aligned}
f_{corr,1D}(x_{sm},d) &= \frac{1}{2\pi\sigma_{sm}\sigma_{seq}} e^{-\left(\frac{x_{sm}^2}{2\sigma_{sm}^2}+\frac{d^2+2dx_{sm}+x_{sm}^2}{2\sigma_{seq}^2}\right)} \\
&= \frac{1}{2\pi\sigma_{sm}\sigma_{seq}} e^{-\frac{x_{sm}^2\sigma_{seq}^2+d^2\sigma_{sm}^2+2dx_{sm}\sigma_{sm}^2+x_{sm}^2\sigma_{sm}^2}{2\sigma_{sm}^2\sigma_{seq}^2}} \\
&= \frac{1}{2\pi\sigma_{sm}\sigma_{seq}} e^{-\frac{(\sigma_{sm}^2+\sigma_{seq}^2)x_{sm}^2+2d\sigma_{sm}^2x_{sm}+d^2\sigma_{sm}^2}{2\sigma_{sm}^2\sigma_{seq}^2}}
\end{aligned}
$$

Now we get all possibilities for a specific d by integrating over $x_{sm}$ from -∞ to ∞:

$$
\begin{aligned}
f_{corr,1D}(d) &= \int_{-\infty}^{\infty} f(x_{sm},d)\,dx_{sm} \\
&= \int_{-\infty}^{\infty} \frac{1}{2\pi\sigma_{sm}\sigma_{seq}} e^{-\frac{(\sigma_{sm}^2+\sigma_{seq}^2)x_{sm}^2+2d\sigma_{sm}^2x_{sm}+d^2\sigma_{sm}^2}{2\sigma_{sm}^2\sigma_{seq}^2}}\,dx_{sm} \\
&= \frac{1}{2\pi\sigma_{sm}\sigma_{seq}} \int_{-\infty}^{\infty} e^{-\frac{(\sigma_{sm}^2+\sigma_{seq}^2)x_{sm}^2+2d\sigma_{sm}^2x_{sm}+d^2\sigma_{sm}^2}{2\sigma_{sm}^2\sigma_{seq}^2}}\,dx_{sm} \\
&= \frac{1}{2\pi\sigma_{sm}\sigma_{seq}} \int_{-\infty}^{\infty} e^{-\left(\frac{\sigma_{sm}^2+\sigma_{seq}^2}{2\sigma_{sm}^2\sigma_{seq}^2}x_{sm}^2+\frac{d}{\sigma_{seq}^2}x_{sm}+\frac{d^2}{2\sigma_{seq}^2}\right)}\,dx_{sm}
\end{aligned}
$$

Using the standard integral

$$
\begin{aligned}
\int_{-\infty}^{\infty} e^{-(ax^2+bx+c)}\,dx &= \sqrt{\frac{\pi}{a}}e^{\left(\frac{b^2-4ac}{4a}\right)} \\
&= \sqrt{\frac{\pi}{a}}e^{\left(\frac{b^2}{4a}-c\right)}
\end{aligned}
$$

with

$$
\begin{aligned}
a &= \frac{\sigma_{sm}^2+\sigma_{seq}^2}{2\sigma_{sm}^2\sigma_{seq}^2} \\
b &= \frac{d}{\sigma_{seq}^2} \\
c &= \frac{d^2}{2\sigma_{seq}^2}
\end{aligned}
$$

we get

$$f_{corr,1D}(d) = \frac{1}{2\pi\sigma_{sm}\sigma_{seq}} \sqrt{2\pi \frac{\sigma_{sm}^2\sigma_{seq}^2}{\sigma_{sm}^2+\sigma_{seq}^2}} e^{\left(\frac{d^2\, 2\sigma_{sm}^2\sigma_{seq}^2}{4\sigma_{seq}^4(\sigma_{sm}^2+\sigma_{seq}^2)} - \frac{d^2}{2\sigma_{seq}^2}\right)}$$

$$= \sqrt{\frac{1}{2\pi(\sigma_{sm}^2+\sigma_{seq}^2)}} e^{\left(\frac{2d^2\sigma_{sm}^2\sigma_{seq}^2}{4\sigma_{seq}^4(\sigma_{sm}^2+\sigma_{seq}^2)} - \frac{d^2}{2\sigma_{seq}^2}\right)}$$

Then with

$$\frac{2d^2\sigma_{sm}^2\sigma_{seq}^2}{4\sigma_{seq}^4(\sigma_{sm}^2+\sigma_{seq}^2)} = \frac{d^2\sigma_{sm}^2}{2\sigma_{seq}^2(\sigma_{sm}^2+\sigma_{seq}^2)} = \frac{d^2}{2\sigma_{seq}^2} - \frac{d^2}{2(\sigma_{sm}^2+\sigma_{seq}^2)}$$

we get

$$f_{corr,1D}(d) = \sqrt{\frac{1}{2\pi(\sigma_{sm}^2+\sigma_{seq}^2)}} e^{\left(\frac{d^2}{2\sigma_{seq}^2} - \frac{d^2}{2(\sigma_{sm}^2+\sigma_{seq}^2)} - \frac{d^2}{2\sigma_{seq}^2}\right)}$$

$$= \frac{1}{\sqrt{2\pi(\sigma_{sm}^2+\sigma_{seq}^2)}} e^{-\frac{d^2}{2(\sigma_{sm}^2+\sigma_{seq}^2)}}$$

$$= \frac{1}{\sqrt{2\pi\sigma^2}} e^{-\frac{d^2}{2\sigma^2}}$$

where we used $\sigma_{sm}^2 + \sigma_{seq}^2 = \sigma^2$.

In two dimensions

$$r = \sqrt{d_x^2 + d_y^2}$$

and the probability of a specific $d_x$ and $d_y$ is:

$$f_{corr}(d_x, d_y) = f_{corr,1D}(d_x) f_{corr,1D}(d_y)$$

$$= \frac{1}{\sqrt{2\pi\sigma^2}} e^{-\frac{d_x^2}{2\sigma^2}} \frac{1}{\sqrt{2\pi\sigma^2}} e^{-\frac{d_y^2}{2\sigma^2}}$$

$$= \frac{1}{2\pi\sigma^2} e^{-\frac{d_x^2+d_y^2}{2\sigma^2}}$$

where we assume $\sigma_x = \sigma_y = \sigma$.

However, multiple combinations of $d_x$ and $d_y$ can lead to the same r. More specifically, we need to integrate over all possible ds. We can determine this by going to polar coordinates

$$
\begin{aligned}
f_{corr}(r) &= \iint_{r^2=d_x^2+d_y^2} \frac{1}{2\pi\sigma^2} e^{-\frac{d_x^2+d_y^2}{2\sigma^2}} dd_x dd_y \\
&= \int_{-\pi}^{\pi} \frac{1}{2\pi\sigma^2} e^{-\frac{r^2}{2\sigma^2}} r d\theta \\
&= \frac{1}{2\pi\sigma^2} e^{-\frac{r^2}{2\sigma^2}} r \int_{-\pi}^{\pi} d\theta \\
&= \frac{1}{2\pi\sigma^2} e^{-\frac{r^2}{2\sigma^2}} 2\pi r \\
&= \frac{r}{\sigma^2} e^{-\frac{r^2}{2\sigma^2}}
\end{aligned}
$$

Note that taking the integral from -π to π, includes the cases where $x_{seq} > x_{sm}$ and $x_{seq} < x_{sm}$, and $y_{seq} > y_{sm}$ and $y_{seq} < y_{sm}$. If we would have included this before, then we would only take the integral from 0 to π/2.

To check if the integral over all probabilities is 1:

$$\int_0^{\infty} f_{corr}(r) dr = 1$$

we use the standard integral

$$\int_0^{\infty} x^n e^{-ax^2} dx = \frac{k!}{2(a^{k+1})} \text{ for } n = 2k+1, k \text{ integer }, a > 0$$

with

$$
\begin{aligned}
n &= 1 \\
k &= \frac{n-1}{2} = 0 \\
a &= \frac{1}{2\sigma^2}
\end{aligned}
$$

to get

$$\int_0^\infty f_{corr}(r)dr = \int_0^\infty \frac{r}{\sigma^2} e^{-\frac{r^2}{2\sigma^2}} dr$$

$$= 2a \int_0^\infty r e^{-ar^2} dr$$

$$= 2a \frac{0!}{2(a^{0+1})} = \frac{2a}{2a} = 1$$

*Cumulative probability function*

To find the cumulative probability function $F_{corr}$ to have the corresponding point within radius r of the initial point, we use the standard integral

$$\int x e^{-cx^2} dx = -\frac{1}{2c} e^{-cx^2}$$

with

$$c = \frac{1}{2\sigma^2}$$

to get

$$F_{corr}(r) = \int_0^r f_{corr}(r')dr'$$

$$= \int_0^r \frac{r'}{\sigma^2} e^{-\frac{r'^2}{2\sigma^2}} dr'$$

$$= 2c \int_0^r r' e^{-cr'^2} dr'$$

$$= 2c \left[ -\frac{1}{2c} e^{-cr'^2} \right]_0^r$$

$$= -\left[ e^{-cr^2} - e^{-c0^2} \right]$$

$$= 1 - e^{-cr^2}$$

$$= 1 - e^{-\frac{r^2}{2\sigma^2}}$$

*Binned probability density*

To obtain the binned probability density for bin width Δr we need to integrate over the bin:

$$
\begin{aligned}
f_{corr,binned}(r,\Delta r) &= \int_{r-\Delta r/2}^{r+\Delta r/2} f_{corr}(r')dr' \\
&= \int_{0}^{r+\Delta r/2} f_{corr}(r')dr' - \int_{0}^{r-\Delta r/2} f_{corr}(r')dr' \\
&= F_{corr}(r+\Delta r/2) - F_{corr}(r-\Delta r/2) \\
&= \left(1-e^{-\frac{(r+\Delta r/2)^2}{2\sigma^2}}\right) - \left(1-e^{-\frac{(r-\Delta r/2)^2}{2\sigma^2}}\right) \\
&= e^{-\frac{r^2-r\Delta r+\Delta r^2/4}{2\sigma^2}} - e^{-\frac{r^2+r\Delta r+\Delta r^2/4}{2\sigma^2}} \\
&= e^{-\frac{r^2+\Delta r^2/4}{2\sigma^2}}\left(e^{\frac{r\Delta r}{2\sigma^2}} - e^{-\frac{r\Delta r}{2\sigma^2}}\right) \\
&= 2e^{-\frac{r^2+\Delta r^2/4}{2\sigma^2}} \sinh\left(\frac{r\Delta r}{2\sigma^2}\right)
\end{aligned}
$$

*Probability*

The probability of finding corresponding points within a distance r is the same as the cumulative probability function:

$$P_{corr}(X<r) = F_{corr}(r) = 1-e^{-\frac{r^2}{2\sigma^2}}$$

*Distance distribution*

To obtain the density of distances between the single-molecule and sequencing datasets, we need to multiply the probability density by the total number of corresponding point pairs:

$$N_{corr} = \alpha_{sm}\alpha_{seq}N$$

The distance distribution functions then become:

$$
\begin{aligned}
d_{corr}(r) &= \alpha_{sm}\alpha_{seq}N\frac{r}{\sigma^2}e^{-\frac{r^2}{2\sigma^2}} \\
d_{corr,binned}(r,\Delta r) &= 2\alpha_{sm}\alpha_{seq}Ne^{-\frac{r^2+\Delta r^2/4}{2\sigma^2}}\sinh\left(\frac{r\Delta r}{2\sigma^2}\right) \\
D_{corr}(r,\Delta r) &= \alpha_{sm}\alpha_{seq}N\left(1-e^{-\frac{r^2}{2\sigma^2}}\right)
\end{aligned}
$$

**Probabilities of true positives, false positives, true negatives, and false negatives**

To determine the probabilities of the true positives (TP), false positives (FP), true negatives (TN) and false negatives (FN), the various cases of occurring points have to be assessed. In total we distinguish 33 cases, each of which we label as TP if the point pair is correctly identified, FP if an erroneous point pair is identified, TN if the point pair is not identified due to one or both of the points being missing, and FN if the point pair is not identified while both points are present (Table S1).

There are four possibilities regarding the presence of the single-molecule (S) and sequencing (D) point pairs:

$$\begin{aligned} P_{corr}(sm, seq) &= \alpha_{sm}\alpha_{seq} \\ P_{corr}(sm, \cancel{seq}) &= \alpha_{sm}(1-\alpha_{seq}) \\ P_{corr}(\cancel{sm}, seq) &= (1-\alpha_{sm})\alpha_{seq} \\ P_{corr}(\cancel{sm}, \cancel{seq}) &= (1-\alpha_{sm})(1-\alpha_{seq}) \end{aligned}$$

In the case that both of the corresponding points are present, the distance between them can be smaller or larger than the threshold R:

$$\begin{aligned} P_{corr}(sm, seq, x < R) &= \alpha_{sm}\alpha_{seq}P_{corr}(x < R) \\ P_{corr}(sm, seq, x > R) &= \alpha_{sm}\alpha_{seq}\left(1 - P_{corr}(x < R)\right) \end{aligned}$$

In addition to considering the corresponding points, in all cases additional random points need to be considered that could appear in the area(s) within the threshold around the corresponding points.

In case both corresponding points are within the distance threshold, no additional random points would result in the point pair to be correctly identified:

$$P_{TP} = \alpha_{sm}\alpha_{seq}P_{corr}(x < R)P_{rand}^{sm}(N = 0)P_{rand}^{seq}(N = 0)$$

If there are random points, then the point pair will be a FN:

$$P_{FN,1} = \alpha_{sm}\alpha_{seq}P_{corr}(x < R)\left(1 - P_{rand}^{sm}(N = 0)P_{rand}^{seq}(N = 0)\right)$$

In case the points of a point pair cannot be detected together because they are separated by a larger distance than the threshold, then we have to consider the possibility that a random point can be detected as a correct match.

In case there is no random point within the vicinity of the original point, or more than one random point, then point pair will be classified as a FN:

$$\begin{aligned} P_{FN,2} &= \frac{1}{2}\alpha_{sm}\alpha_{seq}\left(1-P_{corr}(x<R)\right)\left(1-P_{rand}^{sm}(N=1)\right)+\frac{1}{2}\alpha_{sm}\alpha_{seq}P_{corr}(x>R)\left(1-P_{rand}^{seq}(N=1)\right) \\ &= \frac{1}{2}\alpha_{sm}\alpha_{seq}\left(1-P_{corr}(x<R)\right)\left[\left(1-P_{rand}^{sm}(N=1)\right)+\left(1-P_{rand}^{seq}(N=1)\right)\right] \end{aligned}$$

where the factor ½ to compensate for the fact that both points from the original point pair can be form a new apparently corresponding point pair.

In case we have a sequencing point without a corresponding single-molecule point, then a random single-molecule point can be detected as a false positive point pair. However, the random single-molecule point will also have a corresponding point in the sequencing data. Therefore, there are two situations in which replacement can occur: either the distance between them is larger than the distance threshold R, or the corresponding point is not visible. Additionally, the random point should not have any other random points in the other point set.

$$\begin{aligned} P_{replaced}^{sm} &= P_{rand}^{sm}(N=1)\left[\alpha_{seq}\left(1\text{-}P_{corr}(x<R)\right)+\ (1\text{-}\alpha_{seq})\right]P_{rand}^{seq}(N=0) \\ &= P_{rand}^{sm}(N=1)\left[\alpha_{seq}-\alpha_{seq}P_{corr}(x<R)+\ 1\text{-}\alpha_{seq}\right]P_{rand}^{seq}(N=0) \\ &= P_{rand}^{sm}(N=1)\left[1-\alpha_{seq}P_{corr}(x<R)\right]P_{rand}^{seq}(N=0) \\ P_{replaced}^{seq} &= P_{rand}^{seq}(N=1)\left[\alpha_{sm}\left(1\text{-}P_{corr}(x<R)\right)+(1\text{-}\alpha_{sm})\right]P_{rand}^{sm}(N=0)= \\ &= P_{rand}^{seq}(N=1)\left[1-\alpha_{sm}P_{corr}(x<R)\right]P_{rand}^{sm}(N=0) \end{aligned}$$

If the additional random point has its corresponding point closer than the threshold, or has additional random points within its distance threshold, then the points are not singly matched anymore, and they are excluded resulting in a FN.

$$
\begin{aligned}
P^{sm}_{not-replaced} &= P^{sm}_{rand}(N=1)\left\{\alpha_{seq}\left[P_{corr}(x<R) \;+\; \left(1\text{-}P_{corr}(x<R)\right)\left(1\text{-}P^{seq}_{rand}(N=0)\right)\right]+(1\text{-}\alpha_{seq})\left(1\text{-}P^{seq}_{rand}(N=0)\right)\right\} \\
&= P^{sm}_{rand}(N=1)\left\{\begin{matrix}\alpha_{seq}P_{corr}(x<R) \;+\; \alpha_{seq}\text{-}\alpha_{seq}P_{corr}(x<R)\text{-}\alpha_{seq}P^{seq}_{rand}(N=0)+ \\ \alpha_{seq}P_{corr}(x<R)P^{seq}_{rand}(N=0)+1\text{-}\alpha_{seq}\text{-}P^{seq}_{rand}(N=0)+\alpha_{seq}P^{seq}_{rand}(N=0)\end{matrix}\right\} \\
&= P^{sm}_{rand}(N=1)\left\{1\text{-}P^{seq}_{rand}(N=0)+\alpha_{seq}P_{corr}(x<R)P^{seq}_{rand}(N=0)\right\} \\
&= P^{sm}_{rand}(N=1)\left\{1-\left[1-\alpha_{seq}P_{corr}(x<R)\right]P^{seq}_{rand}(N=0)\right\} \\
&= P^{sm}_{rand}(N=1)\left\{1-P^{sm}_{replaced}\right\} \\
P^{seq}_{not-replaced} &= P^{seq}_{rand}(N=1)\left\{\alpha_{sm}\left[P_{corr}(x<R) \;+\; \left(1\text{-}P_{corr}(x<R)\right)\left(1\text{-}P^{sm}_{rand}(N=0)\right)\right]+(1\text{-}\alpha_{sm})\left(1\text{-}P^{sm}_{rand}(N=0)\right)\right\} \\
&= P^{seq}_{rand}(N=1)\left\{1-P^{seq}_{replaced}\right\}
\end{aligned}
$$

We then get the contributions for FP an FN:

$$
\begin{aligned}
P_{FP,1} &= \frac{1}{2}\alpha_{sm}\alpha_{seq}\left(1-P_{corr}(x<R)\right)\left(P^{sm}_{replaced}+P^{seq}_{replaced}\right) \\
P_{FN,3} &= \frac{1}{2}\alpha_{sm}\alpha_{seq}\left(1-P_{corr}(x<R)\right)\left(\left(1-P^{sm}_{replaced}\right)+\left(1-P^{seq}_{replaced}\right)\right)
\end{aligned}
$$

If only one of the points in a pair is visible, then then without random points or more than one random point, the point is classified as a TN:

$$
P_{TN,1} = \alpha_{sm}(1-\alpha_{seq})\left(1-P^{seq}_{rand}(N=1)\right)+(1-\alpha_{sm})\alpha_{seq}\left(1-P^{sm}_{rand}(N=1)\right)
$$

In addition, the point can again be replaced by a random point, giving similar probability contributions:

$$
\begin{aligned}
P_{FP,2} &= \alpha_{sm}(1-\alpha_{seq})P^{seq}_{replaced}+(1-\alpha_{sm})\alpha_{seq}P^{sm}_{replaced} \\
P_{TN,2} &= \alpha_{sm}(1-\alpha_{seq})\left(1-P^{seq}_{replaced}\right)+(1-\alpha_{sm})\alpha_{seq}\left(1-P^{sm}_{replaced}\right)
\end{aligned}
$$

Finally, there is the case when both the single-molecule and sequencing points from a corresponding pair are not detected, resulting in a TN:

$$
P_{TN,3} = (1-\alpha_{sm})(1-\alpha_{seq})
$$

***Table S1: Overview possibilities when detecting corresponding pairs in a randomized point set.*** *S1 and D1 indicate the corresponding points that are to be detected. TP, FP, TN and FN indicate true* *positive, false positive, true negative and false negative, respectively; the additional number corresponds to the equations above. R indicates the distance threshold. NA means not applicable.*

| | | | S1-D1 | | | | | S2-D2 | | | | |
|---|---|---|---|---|---|---|---|---|---|---|---|---|
| | | | S1 | D1 | Distance | Random D points around S1 | Random S points around D1 | S2 | D2 | Distance | Random D points around S2 | Random S points around D2 |
| \|S1 D1\| | TP | | 1 | 1 | <R | 0 | 0 | | | | | |
| \|S1 D1 S2+\| | FN 1 | | | | | 0 | >0 | | | | | |
| \|S1 D1 D2+\| | FN 1 | | | | | >0 | 0 | | | | | |
| \|S1 D1 S2+ D2+\| | FN 1 | | | | | >0 | >0 | | | | | |
| \|S1\|D1 | FN 2 | ½ | | | >R | 0 | NA | | | | | |
| \|S1 S2 D2\|D1 | FN 3 | ½ | | | | 1 | NA | 1 | NA | <R | NA | NA |
| S2\|S1 D2\|D1 | FP 1 | ½ | | | | | | | | >R | NA | 0 |
| S2\|S1 D2 S3\|D1 | FN 3 | ½ | | | | | | | | | NA | >0 |
| \|S1 D2\|D1 | FP 1 | ½ | | | | | | 0 | | NA | NA | 0 |
| \|S1 D2 S3\|D1 | FN 3 | ½ | | | | | | | | | NA | >0 |
| \|S1 D2 D3+\|D1 | FN 2 | ½ | | | | >1 | NA | | | | | |
| S1\|D1\| | FN 2 | ½ | | | | NA | 0 | | | | | |
| S1\|D1 S2 D2\| | FN 3 | ½ | | | | NA | 1 | NA | 1 | <R | NA | NA |
| S1\|D1 S2\|D2 | FP 1 | ½ | | | | | | | | >R | 0 | NA |
| S1\|D1 S2 D3\|D2 | FN 3 | ½ | | | | | | | | | >0 | NA |
| S1\|D1 S2\| | FP 1 | ½ | | | | | | | 0 | NA | 0 | NA |
| S1\|D1 S2 D3\| | FN 3 | ½ | | | | | | | | | >0 | NA |
| S1\|D1 S2 S3\| | FN 2 | ½ | | | | NA | >1 | | | | | |
| \|S1\| | TN 1 | | 1 | 0 | NA | 0 | NA | | | | | |
| \|S1 S2 D2\| | TN 2 | | | | | 1 | NA | 1 | NA | <R | NA | NA |
| S2\|S1 D2\| | FP 2 | | | | | | | | | >R | NA | 0 |
| S2\|S1 D2 S3\| | TN 2 | | | | | | | | | | NA | >0 |
| \|S1 D2\| | FP 2 | | | | | | | 0 | | NA | NA | 0 |
| \|S1 D2 S3\| | TN 2 | | | | | | | | | | NA | >0 |
| \|S1 D2 D3+\| | TN 1 | | | | | >1 | NA | | | | | |
| \|D1\| | TN 1 | | 0 | 1 | NA | NA | 0 | | | | | |
| \|D1 S2 D2\| | TN 2 | | | | | NA | 1 | NA | 1 | <R | NA | NA |
| \|D1 S2\|D2 | FP 2 | | | | | | | | | >R | 0 | NA |
| \|D1 S2 D3\|D2 | TN 2 | | | | | | | | | | >0 | NA |
| \|D1 S2\| | FP 2 | | | | | | | | 0 | NA | 0 | NA |
| \|D1 S2 D3\| | TN 2 | | | | | | | | | | >0 | NA |
| \|D1 S2 S3+\| | TN 1 | | | | | NA | >1 | | | | | |
| \|\| | TN 3 | | 0 | 0 | NA | NA | NA | | | | | |

The final equations for the TP, FN, FP and TN then become:

$$P_{TP} = \alpha_{sm}\alpha_{seq}P_{corr}(x<R)P_{rand}^{sm}(N=0)P_{rand}^{seq}(N=0)$$

$$P_{FN} = \alpha_{sm}\alpha_{seq}\begin{bmatrix} P_{corr}(x<R)\left(1\text{-}P_{rand}^{sm}(N=0)P_{rand}^{seq}(N=0)\right)+ \\ \frac{1}{2}\left(1-P_{corr}(x<R)\right)\left(4-P_{rand}^{sm}(N=1)-P_{rand}^{seq}(N=1)-P_{replaced}^{sm}-P_{replaced}^{seq}\right) \end{bmatrix}$$

$$P_{FP} = \frac{1}{2}\alpha_{sm}\alpha_{seq}\left(1-P_{corr}(x<R)\right)\left(P_{replaced}^{sm}+P_{replaced}^{seq}\right)+$$

$$\alpha_{sm}(1-\alpha_{seq})P_{replaced}^{seq}+(1-\alpha_{sm})\alpha_{seq}P_{replaced}^{sm}$$

$$P_{TN} = \alpha_{sm}(1-\alpha_{seq})\left(2-P_{rand}^{seq}(N=1)-P_{replaced}^{seq}\right)+$$

$$(1-\alpha_{sm})\alpha_{seq}\left(2-P_{rand}^{sm}(N=1)-P_{replaced}^{sm}\right)+$$

$$(1-\alpha_{sm})(1-\alpha_{seq})$$
